# Ultrafast and Reproducible Proteomics from Small Amounts of Heart Tissue Enabled by Azo and timsTOF Pro

**DOI:** 10.1101/2021.05.24.445470

**Authors:** Timothy J. Aballo, David S. Roberts, Jake A. Melby, Kevin M. Buck, Kyle A. Brown, Ying Ge

**Affiliations:** Molecular and Cellular Pharmacology Training Program, University of Wisconsin-Madison, Madison, Wisconsin 53705, USA; Department of Cell and Regenerative Biology, University of Wisconsin-Madison, Madison, Wisconsin 53705, USA; Department of Chemistry, University of Wisconsin-Madison, Madison, Wisconsin 53706, USA; Department of Surgery, University of Wisconsin-Madison, Madison, Wisconsin, 53706, USA; Human Proteomics Program, School of Medicine and Public Health, University of Wisconsin-Madison, Madison, Wisconsin 53705, USA

**Keywords:** photocleavable surfactant, human heart proteomics, quantitative proteomics, sample preparation, bottom-up proteomics

## Abstract

Global bottom-up mass spectrometry (MS)-based proteomics is widely used for protein identification and quantification to achieve a comprehensive understanding of the composition, structure, and function of the proteome. However, traditional sample preparation methods are time-consuming, typically including overnight tryptic digestion, extensive sample clean-up to remove MS-incompatible surfactants, and offline sample fractionation to reduce proteome complexity prior to online liquid chromatography-tandem mass spectrometry (LC-MS/MS) analysis. Thus, there is a need for a fast, robust, and reproducible method for protein identification and quantification from complex proteomes. Herein, we developed an ultrafast bottom-up proteomics method enabled by Azo, a photocleavable, MS-compatible surfactant that effectively solubilizes proteins and promotes rapid tryptic digestion, combined with the Bruker timsTOF Pro, which enables deeper proteome coverage through trapped ion mobility spectrometry (TIMS) and parallel accumulation-serial fragmentation (PASEF) of peptides. We applied this method to analyze the complex human cardiac proteome and identified nearly 4,000 protein groups from as little as 1 mg of human heart tissue in a single one-dimensional LC-TIMS-MS/MS run with high reproducibility. Overall, we anticipate this ultrafast, robust, and reproducible bottom-up method empowered by both Azo and the timsTOF Pro will be generally applicable and greatly accelerate the throughput of large-scale quantitative proteomic studies. Raw data are available via the MassIVE repository with identifier MSV000087476.

## Introduction

Global bottom-up mass spectrometry (MS)-based proteomics is widely used for protein identification and quantification to achieve a comprehensive understanding of the composition, structure, and function of the proteome.^1-3^ However, typical bottom-up methods are time-consuming, involving extraction and solubilization of proteins, reduction and alkylation of disulfide bonds, enzymatic digestion of proteins into peptides, and removal of MS-incompatible surfactants prior to online liquid chromatography (LC)-tandem MS (MS/MS).^3-7^ Surfactants are commonly used during sample preparation to improve protein solubility, especially for membrane proteins; however, most surfactants are MS-incompatible and thus need to be removed before LC-MS/MS which further complicates and extends the sample clean-up process.^7-12^

Recently, we developed an anionic, photocleavable surfactant, 4-hexylphenylazosulfonate (referred to as “Azo”), which promotes efficient extraction and solubilization of proteins with similar performance to SDS but can rapidly degrade upon UV exposure.^13, 14^ Importantly, we have shown the unique capability of Azo as an “all-in-one” MS-compatible surfactant for both top-down and bottom-up proteomics, which enables high-throughput sample handling before MS-analysis.^13^ Azo promotes rapid and robust digestion, even for trypsin-resistant proteins, and thus Azo-aided bottom-up methods have been successfully developed for effective membrane and extracellular matrix proteomics.13, 15 As enzymatic digestion is a major time-consuming step in bottom-up sample preparation, removing this bottleneck greatly speeds up sample processing time. Nevertheless, the proteome coverage was limited in these Azo-enabled bottom-up proteomics studies using one-dimensional (1D) reversed-phase (RP)LC-MS/MS, in part due to the instrumentation (a standard quadrupole time of flight (Q-TOF) mass spectrometer) used for data acquisition.^13, 15^

To obtain deeper proteome coverage, multidimensional separation (often involving offline sample fractionation) is employed for the analysis of complex proteomes, which adds another time-consuming step in the bottom-up proteomics workflow.^16-20^ Though there have been developments in online multidimensional LC separation coupled to MS analysis, these current online two-dimensional (2D) LC-MS/MS methods remain technically challenging to implement and can lead to decreased sensitivity.^21-23^ Recently, Bruker interfaced trapped ion mobility spectrometry (TIMS) with the fast scan speeds of Q-TOF mass spectrometers in their timsTOF Pro, an instrument that provides an additional dimension of peptide separation in the gas phase. When the timsTOF Pro is operated in parallel accumulation-serial fragmentation (PASEF) mode, peptide ions that co-elute during LC are accumulated in parallel at specific mobility spaces, and then, in conjunction with rapid quadrupole switching, are serially ejected from the TIMS cell for MS/MS analysis.^24, 25^ This process increases sequencing speeds, without a loss in sensitivity, leading to increased proteome sequencing depth.^24, 25^

Here, for the first time, we report an ultrafast and reproducible quantitative global bottom-up proteomics method with broad proteome coverage enabled by Azo and the timsTOF Pro. We applied this method to identify and reproducibly quantify the proteome from very small amounts of heart tissue. We achieved rapid tryptic digestion in 30 min and identified nearly 4,000 protein groups in a single 1D LC-TIMS-MS/MS run from just 1 mg of human left ventricular (LV) tissue. We envision this ultrafast, robust, reproducible, and quantitative bottom-up method can be generally applicable and can greatly accelerate the throughput of large-scale quantitative proteomic studies even when samples are limited.

## Materials and Methods

### Chemicals and reagents

4-hexylphenylazosulfonate (Azo) was synthesized in-house as previously described.^13, 14^ All reagents were purchased from Millipore Sigma (St. Louis, MO, USA) and Fisher Scientific (Fair Lawn, NJ, USA) unless noted otherwise. All solutions were prepared with HPLC-grade water (Fisher Scientific).

### Sample Preparation

Donor hearts were obtained from the University of Wisconsin Organ Procurement Organization. Tissue collection procedures were approved by Institutional Review Board of the University of Wisconsin-Madison (Study # 2013-1264). Donor heart tissue was maintained in cardioplegic solution before dissection, and after dissection, the tissue was immediately snap-frozen in liquid nitrogen and stored at –80 °C.

In a cold room (4 °C), human LV tissue was quickly cut into 1 mg, 5 mg and 20 mg pieces then washed twice in 20 tissue volumes (20 µL/mg) Mg^2+^/Ca^2+^-free DPBS containing 1X HALT protease and phosphatase inhibitor (Thermo Fisher Scientific, Waltham, MA, USA). After washing, tissues were homogenized in 10 tissue volumes (10 µL/mg) of Azo lysis buffer (0.2% w/v Azo, 25 mM ammonium bicarbonate, 10 mM L-methionine, 1 mM dithiothreitol (DTT), and 1X HALT protease and phosphatase) using a Teflon pestle (1.5 mL micro-centrifuge tube, flat tip; Thomas Scientific, Swedesboro, NJ, USA).^13^ Samples were diluted to 0.1% Azo and then homogenized. Samples were centrifuged at 21,100 *g* for 30 min at 4 °C, and the supernatant was transferred to a new tube.

Samples were normalized to 1 mg/mL protein in 0.1% Azo by the Bradford assay (Bio-Rad, Hercules, CA, USA, Cat# 5000006) using bovine serum albumin to generate a standard curve. Disulfide bonds were reduced with 30 mM DTT at 37 °C for 1 h and alkylated with 30 mM iodoacetamide in the dark for 45 min. Trypsin Gold (Promega, Madison, WI, USA) was added in a protein:protease ratio (wt/wt) of 50:1 and incubated for 30 min, 1 h, or 24 h at 37 °C. Enzymatic activity was quenched by acidifying samples to pH 2 with formic acid (FA) and placing samples on ice. After digestion, Azo was degraded at 305 nm (UVN-57 Handheld UV Lamp; Analytik Jena, Jena, TH, DEU) for 5 min, samples were spun at 21,100 *g* at 4 °C for 15 min, and then desalted using 100 µL Pierce C18 tips (Thermo Fisher Scientific) following the manufacturer’s specifications to remove Azo degradants and other salts typically present in the digestion. Desalted peptides were dried in a vacuum centrifuge and reconstituted in 0.1% FA. The peptide concentration was determined by A205 readings using a NanoDrop. Samples were centrifuged at 21,100 *g* at 4 °C for 30 min prior to analysis.

### Bottom-up Data Acquisition

LC-TIMS-MS/MS was carried out using a nanoElute nano-flow ultra-high pressure LC system (Bruker Daltonics, Bremen, Germany) coupled to the timsTOF Pro, a TIMS Q-TOF mass spectrometer (Bruker Daltonics), using a CaptiveSpray nano-electrospray ion source (Bruker Daltonics). For most analyses, 200 ng of peptide digest was loaded on a capillary C18 column (25 cm length, 75 µm inner diameter, 1.6 µm particle size, 120 Å pore size; IonOpticks, Fitzroy, VIC, AUS). To assess instrument sensitivity with complex heart proteome digest, peptide loading amount was decreased from 200 ng per injection down to 6.25 ng. Peptides were separated at 55 °C using a 120 min gradient at a flow rate of 400 nL/min (mobile phase A (MPA): 0.1% FA; mobile phase B (MPB): 0.1% FA in acetonitrile). A stepwise gradient of 2-17% MPB was applied for 60 min, followed by a step from 17-25% MPB from 60-90 min, 25-37% MPB from 90-100 min, 37-85% MPB from 100-110 min, and finished with a wash at 85% MPB for an additional 10 min for a total runtime of 120 min per analysis.

The timsTOF Pro was operated in PASEF mode.^24, 25^ Mass spectra for MS and MS/MS scans were recorded between 100-1700 *m/z*. Ion mobility resolution was set to 0.60-1.60 V*s/cm over a ramp time of 100 ms. Data-dependent acquisition was performed using 10 PASEF MS/MS scans per cycle with a near 100% duty cycle. A polygon filter was applied in the *m/z* and ion mobility space to exclude low *m/z*, singly charged ions from PASEF precursor selection. An active exclusion time of 0.4 min was applied to precursors that reached 20,000 intensity units. Collisional energy was ramped stepwise as a function of ion mobility.^24^ Peptide loading was normalized to the total ion chromatograms from 200 ng injections of K562 whole cell lysate standard (Promega).

### Data Analysis

MS raw files were processed with MaxQuant^26^ (version 1.6.17.0), MS/MS spectra were matched to *in silico* derived tryptic peptide fragment mass values from the Uniprot human database UP000005640 (accessed 04 September 2020) and potential contaminants by the Andromeda search engine.^27^ Peptide mass was limited to 8,000 Da with a maximum of two missed cleavages and a minimum length of 7 amino acids. Methionine oxidation and acetylation of protein N-termini were set as variable modifications, whereas carbamidomethylation of cysteine was set as a fixed modification. Initial maximum mass tolerance was set at 70 ppm for precursor ions and 40 ppm for fragment ions similarly as reported previously.^28^ Individual precursor mass tolerances were applied to each peptide spectral match (PSM) by using the default parameters on MaxQuant. Match between runs was enabled with a retention time matching window of 0.7 min and an ion mobility matching window of 0.05 V*s/cm^2^. Label-free quantification (LFQ) was performed using classic normalization and a minimum ratio count of 2. A reverse sequence library was generated to control the false discovery rate at less than 1% for PSMs and protein group identifications.

After MaxQuant processing, data were further analyzed and visualized using Perseus (version 1.6.14.0).^29^ Data were filtered to remove potential contaminants, reverse hits, and protein groups “only identified by site”. Protein intensities were Log2 transformed, and scatterplots of protein group LFQ intensities were generated between samples and annotated with Pearson correlation coefficients. Comparisons of unique protein groups and unique peptides identified by MaxQuant were visualized with Bio Venn.^30^

## Results and Discussion

### Azo Enabled Ultrafast Proteomics using the timsTOF Pro

The goal of this study was to develop an ultrafast and reproducible quantitative proteomics method from small amounts of heart tissue using the timsTOF Pro (Figure 1A). Previously we demonstrated that Azo, an in-house developed photocleavable and MS compatible anionic surfactant, effectively solubilized proteins and promoted rapid enzymatic digestion.^13^ However, the reproducibility of these ultrafast digestions for quantitative proteomics had not been demonstrated using the timsTOF Pro for data acquisition. As the heart proteome is highly complex and the timsTOF Pro’s enhanced sensitivity enables deep proteome coverage, we set out to test if Azo in combination with the timsTOF Pro could enable ultrafast quantitative proteomics with high reproducibility and high proteome coverage. The timsTOF Pro greatly increases MS/MS sequencing speed through PASEF without sacrificing sensitivity,^24, 25^ a mode in which ions that co-elute from the LC column are accumulated in parallel in the TIMS cell, separated into discreet packets based on their TIMS mobility, and then, in conjunction with rapid quadrupole switching, are serially ejected and subjected to MS/MS (Figure 1B). This process substantially increases the number of protein groups and peptides identified from K562 whole cell lysate digest as a reference standard (Supplementary Figure S1). Utilizing PASEF, we were able to identify 6,364 protein groups and 60,159 peptides from three injections of 200 ng of K562 whole cell lysate, compared to only 2,531 protein groups and 15,815 peptides from the same sample with PASEF and TIMS disabled, exemplifying how PASEF greatly expands proteome coverage.

**Figure 1.**
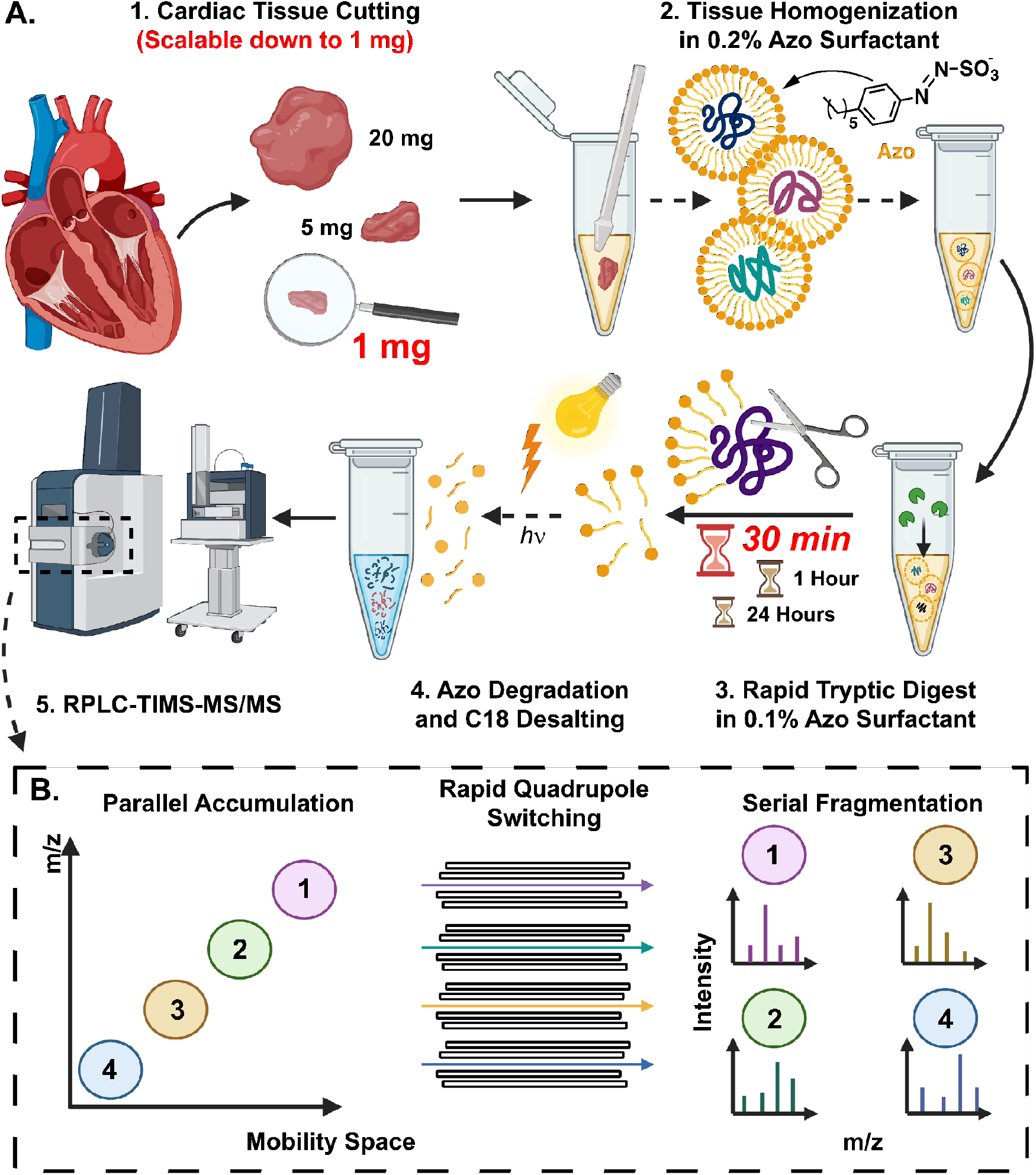
Global bottom-up proteomics workflow using Azo and PASEF on the timsTOF Pro. **(A)** Sample preparation, including (1) tissue cutting (scalable down to 1 mg of tissue), (2) tissue homogenization in Azo, a photocleavable surfactant, (3) rapid tryptic digest (as quick as 30 min), and (4) Azo degradation and C18 desalting prior to (5) RPLC-TIMS-MS/MS. **(B)** Schematic of PASEF on the timsTOF Pro, demonstrating parallel accumulation of ions, rapid quadrupole mass filter switching, and serial ion fragmentation.

Using this platform, we first set out to determine if Azo could enable ultrafast enzymatic digestion of the human heart proteome. Previously, we had shown that Azo could be employed to reduce digestion times down to 30 min, but the proteome coverage in this previous study was limited as data acquisition was performed on the Impact II (Bruker), a standard Q-TOF MS, and without any offline sample fractionation.^13^ Therefore, we sought to evaluate if these ultrafast digestions demonstrate reproducibility for quantitative proteomics on the timsTOF Pro, a sensitive instrument that provides much deeper proteome coverage. To do so, we homogenized three replicates of 20 mg of human LV cardiac tissue, a commonly used tissue for heart proteomics, in an Azo-containing buffer to extract and solubilize cardiac proteins. After sample normalization, thiol-containing cysteine residues were reduced and alkylated and then proteins were digested at 37 °C with trypsin in a 0.1% (w/v) Azo solution as described previously.^13^ Tryptic digestions were halted after 30 min, 1 h, and 24 h and then prepared for LC-TIMS-MS/MS analysis (Figure 1A). After just 30 min of enzymatic digestion in 0.1% Azo, the extracted cardiac proteome was nearly completely digested, similar to 24 h digested samples (Figure 2A). From the data collected by LC-TIMS-MS/MS, we observed a high degree of overlap in protein groups and peptides identified among the different digest times (Figure 2B and 2C, Supplementary Table S1). Notably, within three extraction replicates, 4,016 unique protein groups from 36,312 unique peptides were identified after only 30 min of tryptic digest (Figure 2B and 2C). From these same three extractions, 3,834 protein groups and 37,604 peptides were identified after 1 h of digestion, and 3,841 protein groups and 37,054 peptides were identified after 24 h of digestion (Figures 2B and 2C). These data indicate that 30 min of enzymatic digestion in 0.1% Azo sufficiently digests the cardiac proteome, leading to a similar number of protein group and peptide identifications.

**Figure 2.**
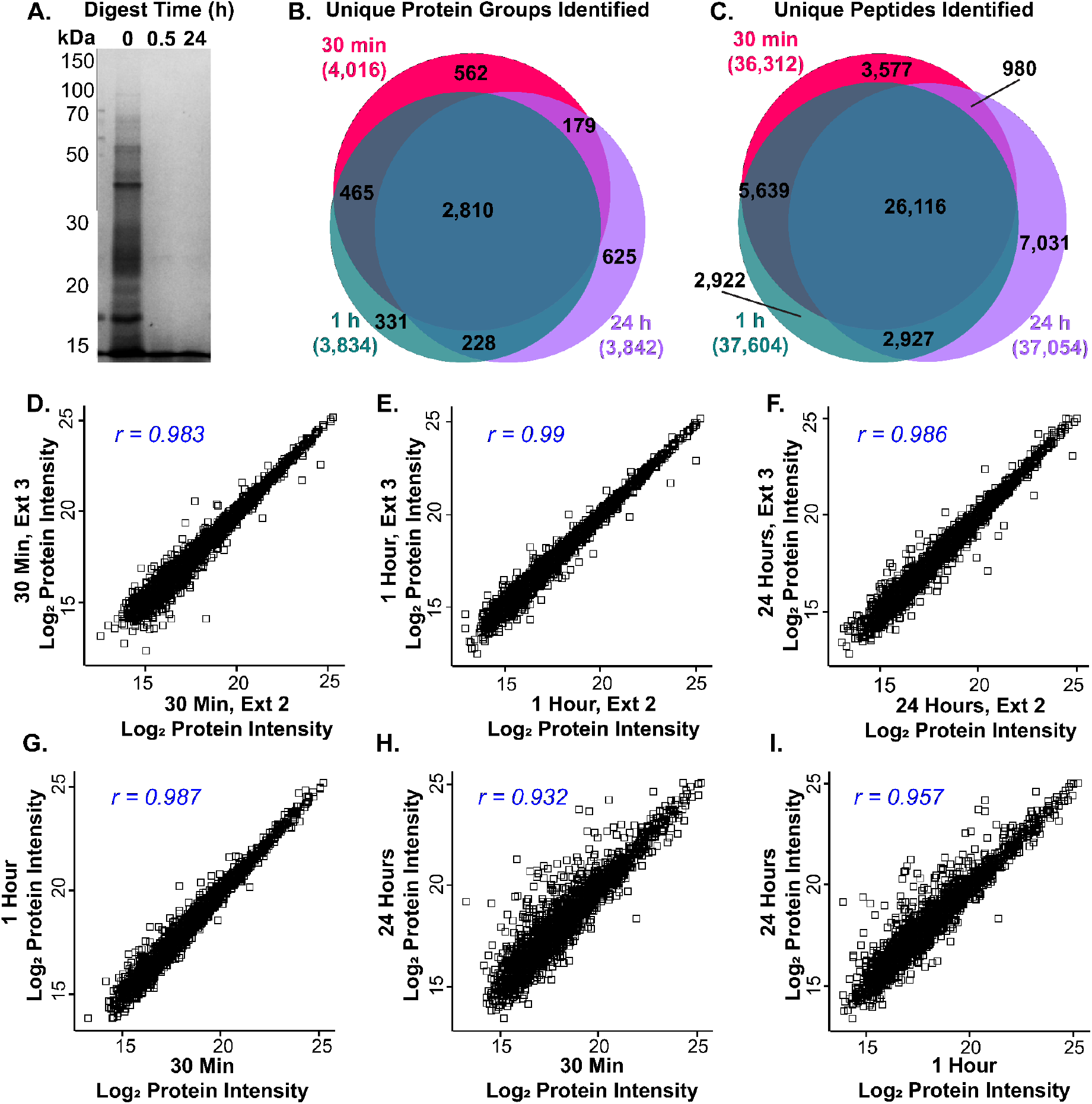
Azo promotes rapid and reproducible tryptic digestion for quantitative proteomics. (**A)** SDS-PAGE showing tryptic digestion efficiency in 0.1% Azo after 0.5 h and 24 h of digestion. **(B and C)** Venn diagrams illustrating overlap in unique protein groups **(B)**, and unique peptides **(C)** identified by MaxQuant between 30 min, 1 h, and 24 h tryptic digestions (n=3 replicates for each group). **(D-F)** Scatterplots of Log2 LFQ protein intensities showing high reproducibility between replicates from 30 min **(D)**, 1 h **(E)**, and 24 h **(F)** tryptic digestions. Pearson correlation coefficients are shown in the top left corner of each panel. (**G-I)** Scatterplots of Log2 LFQ protein intensities showing high reproducibility between averaged replicates from 30 min digestions plotted against 1 h digestions **(G)**, 30 min digestions plotted against 24 h digestions **(H)**, and 1 h digestions plotted against 24 h digestions **(I)** with Pearson correlation coefficients shown in the top left corner of each panel (n=3 replicates for each group).

Subsequently, we sought to evaluate the reproducibility of these ultrafast 30 min tryptic digestions, as reproducible sample preparation and quantitation is critical for future biological studies. Extraction replicates from the same piece of tissue after 30 min tryptic digestions were normally distributed and displayed a strong correlation in protein group LFQ intensities (Figure 2D, Supplementary Figures S2 and S3). These data were similar after 1 h and 24 h digestions (Figure 2E and 2F, Supplementary Figure S2 and S3), suggesting that 30 min rapid digestion lead to highly reproducible and quantitatively similar protein group and peptide LFQ intensities. Furthermore, protein group LFQ intensities between the different digestion times also displayed similar correlations (Figure 2G-I), indicating that increased digest time does not strongly alter protein group LFQ intensity (Supplementary Figure S4). Our data indicates that with 30 min of Azo-enhanced tryptic digest, proteins are well digested, producing similar protein group and peptide LFQ intensities when compared to 1 h and 24 h tryptic digestions.

### Reproducible Proteome Analysis from Small Amounts of Tissue

Next, we sought to determine whether this Azo-enabled sample preparation, including ultrafast tryptic digestion, could be applied to small amounts of cardiac tissue. In studies involving limited amounts of tissue, such as those analyzing precious clinical samples or cardiac tissue harvested from small organisms, reducing the amount of tissue required for protein extraction and processing is highly desirable. Using our method, we were able to extract and analyze a similar number of protein groups and peptides from 20 mg, 5 mg and 1 mg tissue extractions, with 3,047 shared protein groups and 28,945 shared peptides identified among the three tissue extractions (Figure 3A and 3B, Supplementary Figure S5, Supplementary Table S2). Notably, within three extraction replicates from only 1 mg of tissue, we identified 4,114 unique protein groups and 38,319 unique peptides (Figure 3A and 3B). We observed a similar number of unique protein groups and peptides from 20 mg (4,279 and 40,276, respectively) and 5 mg (4,049 and 37,367, respectively) tissue extractions.

**Figure 3.**
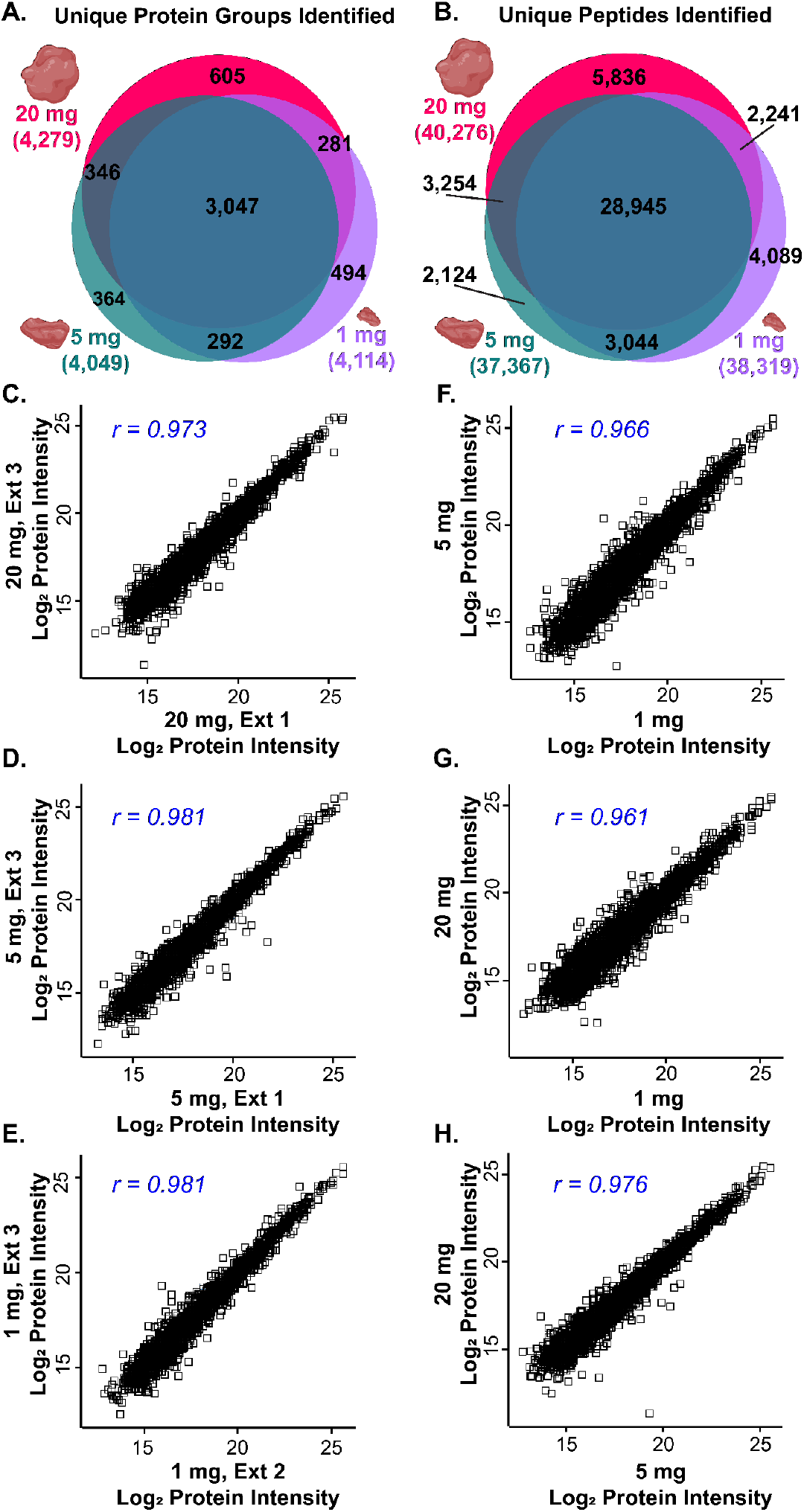
Reproducible protein identification and quantitation from small amounts of tissue. **(A and B)** Venn diagrams illustrating overlap in unique protein groups **(A)**, and unique peptides **(B)** identified by MaxQuant between 20 mg, 5 mg, and 1 mg of tissue (n=3 replicates for each group). **(D-F)**. Scatterplots of Log2 LFQ protein intensities showing high reproducibility between replicates from 20 mg **(D)**, 5 mg **(E)**, and 1 mg **(F)** tissue extractions. Pearson correlation coefficients are shown in the top left corner of each panel. (**G-I)** Scatterplots of Log2 LFQ protein intensities showing high reproducibility between averaged replicates from 1 mg extractions plotted against 5 mg extractions **(G)**, 1 mg extractions plotted against 20 mg extractions **(H)**, and 5 mg extractions plotted against 20 mg extractions **(I)** with Pearson correlation coefficients shown in the top left corner of each panel (n=3 replicates for each group).

As the heart proteome is highly complex, many previous global bottom-up heart proteomics studies employed offline peptide fractionation prior to online LC-MS/MS to increase proteome sequencing depth.^31-37^ For example, a global bottom-up proteomics analysis of heart chamber tissue from rhesus monkeys made use of overnight tryptic digestion, isobaric peptide labelling, and offline sample fractionation before analysis with a highly sensitive Orbitrap Fusion (Thermo), which altogether lead to the identification of 1,633 protein groups.^36^ Similarly, a global proteomic profiling of human LV tissue from patients with end-stage dilated cardiomyopathy identified a total of 4,263 protein groups after 15 h of digestion with Lys-C and trypsin, Sep-Pak desalting, isobaric iTRAQ labeling, and offline pH-based peptide prefractionation prior to online LC-MS/MS.^37^ Moreover, another study by the Mann group identified over 7,000 protein groups from human LV tissue, this was accomplished through extensive offline pH-based fractionation with an automated “loss-less” nano-fractionator,^38^ resulting in 54 total fractions yielding 8 LC-MS/MS runs per sample using Q Exactive HF (Thermo).^31^ Here, using our Azo-enabled bottom-up proteomics method, we identified nearly 4,000 protein groups in a single 1D LC-TIMS-MS/MS run (Supplementary Figure S6), greatly speeding up sample processing and data acquisition while providing deep proteome coverage. Importantly, our method is high-throughput (30 min total digestion time) and does not require extensive sample preparation, which makes this Azo-enabled method more amenable to large-scale biological studies.

Additionally, extraction replicates from 20 mg, 5 mg, and 1 mg of tissue demonstrated highly reproducible, normally distributed protein group LFQ intensities (Figure 3C-E, Supplementary Figure S7 and S8), and protein group LFQ intensities were similar among the different tissue extraction amounts (Figure 3F-H). While there were some differentially extracted protein groups (p≤0.01) among the various tissue amounts (Supplementary Figure S9), these changes are likely due to biological heterogeneity at small scales. Overall, these results indicate that we have developed a robust bottom-up sample processing method that employs ultrafast enzymatic digestion of the human cardiac proteome using Azo and that this method can be scaled down to small tissue amounts while maintaining high proteome coverage and strong reproducibility between extraction replicates for quantitative proteomics.

### High Proteome Coverage from Low Amounts of Peptide on the timsTOF Pro

Lastly, we wanted to investigate the sensitivity and reproducibility of the timsTOF Pro when analyzing low amounts of tryptic peptides. From three different extraction replicates from 20 mg of cardiac tissue, we analyzed 200, 100, 50, 25, 12.5 and 6.25 ng of digested peptides by LC-TIMS-MS/MS. From three injections of just 6.25 ng of peptide, we were able to identify 1,532 unique protein groups and 9,503 unique peptides (Figure 4A and 4B, Supplementary Table S3). While this is 32-fold less peptide mass than loaded in 200 ng injections, we identified only 3-fold more protein groups (4,303) and 4-fold more peptides (40,276) in three 200 ng injections, highlighting the timsTOF Pro’s remarkable sensitivity. Notably, throughout this injection series, protein group LFQ intensities were normally distributed, and there were strong correlations among 200 ng injections and among 6.25 ng injections, indicating there is strong reproducibility even at low peptide loading amounts (Figure 4C and 4D, Supplementary Figures S10 and S11). In cases where sample amounts are extremely limited, the timsTOF Pro, with its enhanced sensitivity due to its TIMS cell, can reproducibly identify a substantial number of protein groups and peptides from very small amounts of tryptic digest.

**Figure 4.**
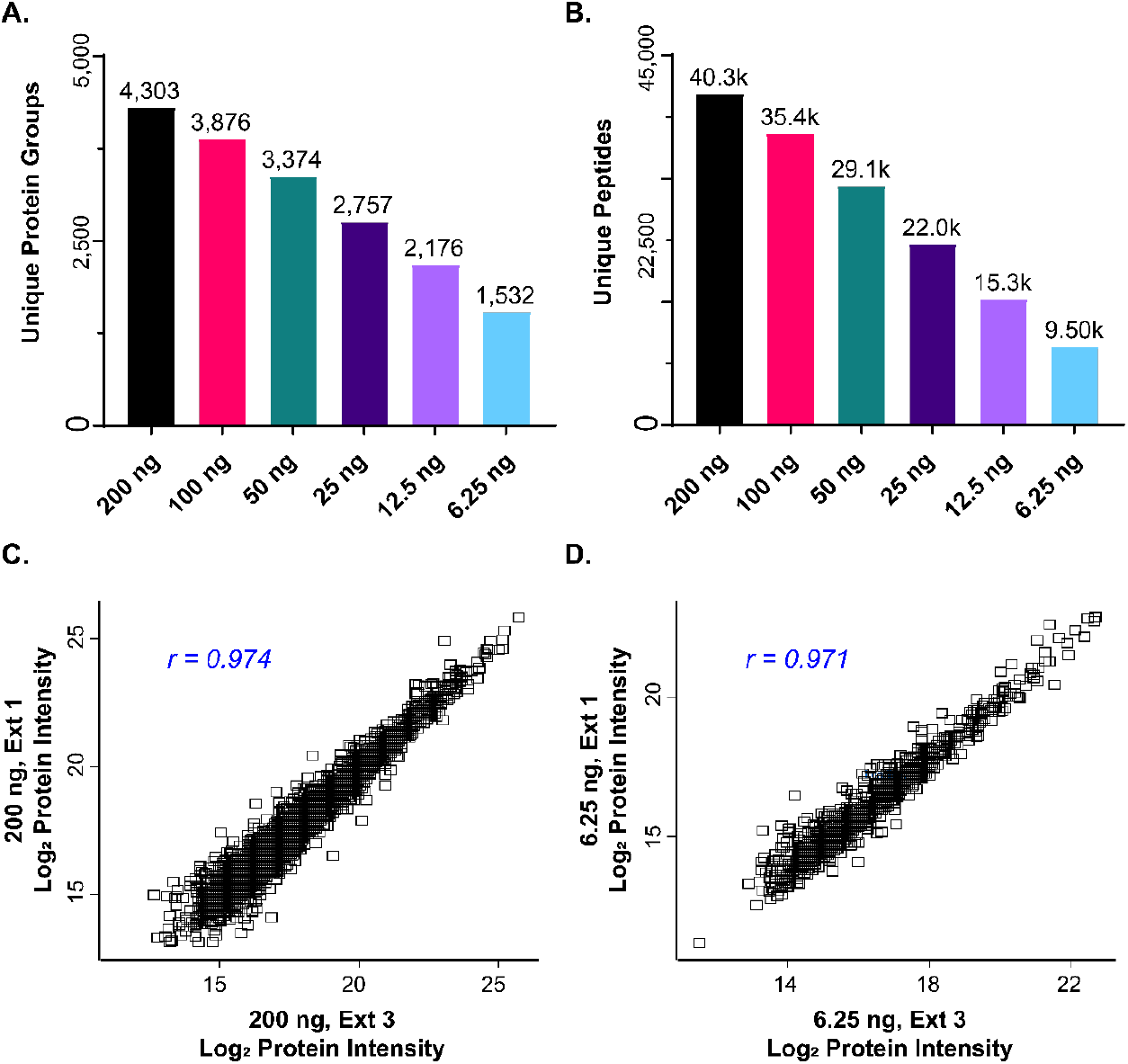
LFQ analysis with high reproducibility from low sample loading amounts on the timsTOF Pro. **(A and B)**. Bar charts showing the total number of unique protein groups **(A)**, and unique peptides (**B)**, identified by MaxQuant from injections of 200, 100, 50, 25, 12.5, and 6.25 ng of digested peptide (n=3 for each group), respectively. Total number of identifications are displayed above each bar. **(C and D)**. Scatterplots of Log2 LFQ protein intensities showing high reproducibility between replicates from 200 ng **(C)** and 6.25 ng **(D)** injections. Pearson correlation coefficients are shown in the top left corner of each panel.

In summary, we have developed an ultrafast bottom-up method for quantitatively analyzing the human cardiac proteome. Our workflow is quick, employing rapid (30 min) tryptic digestion without sacrificing proteome coverage or reproducibility and can be easily scaled down to small amounts of tissue. We achieve deep proteome coverage, identifying nearly 4,000 protein groups in a single LC-TIMS-MS/MS run from a 1 mg tissue extraction without extensive sample clean-up and fractionation, benefiting from both Azo and the timsTOF Pro. Overall, we anticipate this robust, ultrafast, and reproducible bottom-up method for quantitatively analyzing the human cardiac proteome will be the standard procedure for large-scale proteomic studies to answer an array of biological questions not only in cardiac tissue but also in other tissues with highly complex proteomes.

## Associated Content

### Supporting Information

The Supporting Information is available free of charge at

Supplementary Methods:

SDS-Polyacrylamide Gel Electrophoresis (SDS-PAGE)

Bottom-Up Data Acquisition without TIMS/PASEF

Data Analysis for K562 Whole-Cell Digest Comparison

#### Supplementary Figures

Supplementary Figure S1. timsTOF Pro and PASEF enable deep proteome coverage

Supplementary Figure S2. Log2 LFQ protein group intensities for each digestion time are normally distributed.

Supplementary Figure S3. Individual digestion replicates show reproducible Log2 LFQ protein intensities

Supplementary Figure S4. Comparison of protein abundance between different tryptic digestion times

Supplementary Figure S5. SDS-PAGE demonstrating reproducible proteome extraction from different tissue extraction amounts

Supplementary Figure S6. Comparison of protein groups identified between different tissue extraction replicates

Supplementary Figure S7. Log2 LFQ protein group intensities for each tissue extraction amount are normally distributed

Supplementary Figure S8. Individual replicates from different tissue amounts show reproducible Log2 LFQ protein intensities

Supplementary Figure S9. Comparison of proteins extracted between different tissue amounts

Supplementary Figure S10. Log2 LFQ protein group intensities are normally distributed at low peptide loading amounts

Supplementary Figure S11. 200 ng and 6.25 ng peptide injection replicates show reproducible Log2 LFQ protein intensities

#### Supplementary Tables

Supplementary Table S1. Protein groups identified after 24 h, 1 h, and 30 min tryptic digestions

Supplementary Table S2. Protein groups identified from 20 mg, 5 mg, and 1 mg tissue extractions

Supplementary Table S3. Protein groups identified from 200, 100, 50, 25, 12.5, and 6.25 ng peptide injections

## Acknowledgements

Financial support was provided by the National Institutes of Health (NIH) R01 HL096971 (to Y.G.). Y.G. would also like to acknowledge R01 GM117058, GM125085, HL109810 and S10 OD018475. T.JA. would like to acknowledge support from the Training Program in Molecular and Cellular Pharmacology, T32 GM008688-20. J.A.M. would like to acknowledge support from the Training Program in Translational Cardiovascular Science, T32 HL007936-20. K.A.B. would like to acknowledge the Cardiovascular Research Center Training Program in Translational Cardiovascular Science, T32 HL007936-19 and Vascular Surgery Research Training Program grant T32HL110853. We would like to thank Mel Park, Oliver Raether, Guillaume Tremintin, Yue Ju, Conor Mullins, Michael Greig, Gary Kruppa, Paul Speir and Rohan Thakur of Bruker Daltonics for their kind help and provision of the Bruker timsTOF Pro used in this work. Moreover, we would like to acknowledge James Anderson and Carrie Sparks at the University of Wisconsin Organ and Tissue Donation for providing donor hearts. Some aspects of the TOC, Figure 1, and Figure 3 were created using BioRender.com.

## Abbreviations

MS: mass spectrometry
LC: liquid chromatography
MS/MS: tandem MS
1D: one-dimensional
RP: reversed-phase
Q-TOF: quadrupole time of flight
2D: two-dimensional
TIMS: trapped ion mobility
PASEF: parallel-accumulation-serial fragmentation
LV: left ventricular
DTT: dithiothreitol
FA: formic acid
MPA: mobile phase A
MPB: mobile phase B
PSM: peptide spectral match
LFQ: label-free quantification

## TOC

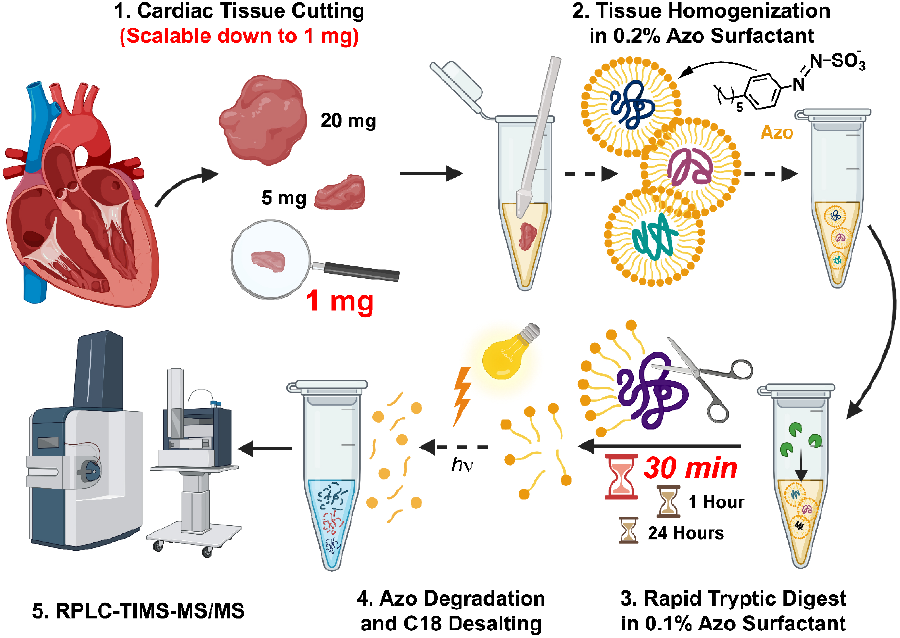

