## Supporting Information for "Ultrafast and Reproducible Proteomics from Small Amounts of Heart Tissue Enabled by Azo and timsTOF Pro"

**Table of Contents**

Supplementary Methods……………………………….……………….……………..…...S-3

Supplementary Figures

Supplementary Figure S1. timsTOF Pro and PASEF enable deep proteome coverage.……………………………….…………………….......………….S-5

Supplementary Figure S2. $Log_2$ LFQ protein group intensities for each digestion time are normally distributed.………………………………………….……..……..…S-6

Supplementary Figure S3. Individual digestion replicates show reproducible $Log_2$ LFQ protein intensities……………………………………………………….….…S-7

Supplementary Figure S4. Comparison of protein abundance between different tryptic digestion times...……………………..……………………………….…….…S-8

Supplementary Figure S5. SDS-PAGE demonstrating reproducible proteome extraction from different tissue amounts…………..………………………………..……..…..S-9

Supplementary Figure S6. Comparison of protein groups identified between different tissue extraction replicates………………………………………………………..S-10

Supplementary Figure S7. $Log_2$ LFQ protein group intensities for each tissue amount are normally distributed……….……………………….......................................S-11

Supplementary Figure S8. Individual replicates from different tissue amounts show reproducible $Log_2$ LFQ protein intensities…………….………..……...………..…S-12

Supplementary Figure S9. Comparison of proteins extracted between different tissue amounts.........................................................................................................S-13

Supplementary Figure S10. $Log_2$ LFQ protein group intensities are normally distributed at low peptide loading amounts……………...……………....…………………..…S-14

Supplementary Figure S11. 200 ng and 6.25 ng peptide injection replicates show reproducible $Log_2$ LFQ protein intensities.………………...………………...….……S-15

References………………………………………………………………………..S-16

### Supplementary Methods

**SDS-Polyacrylamide Gel Electrophoresis (SDS-PAGE)**. For SDS-PAGE analysis to evaluate tryptic digestion times, samples were diluted with an equal volume of 2X Laemmli Sample Buffer. 7.5 µL PageRuler™ unstained protein ladder (Thermo Scientific, Waltham, MA) or 7.5 µg total protein per sample was loaded into individual wells of a hand-cast 12.5% polyacrylamide SDS-PAGE gel (15 comb well, 10.0 cm × 10.0 cm, 1.0 mm thick). Samples were electrophoretically migrated through the stacking gel at 70 V for 30 min, then through the resolving gel at 150V for 90 min. The gel was washed briefly with nanopure water, stained with a Coomassie Brilliant Blue (CBB) solution (500 mg CBB R-250, 10% acetic acid, 50% methanol in nanopure water) and destained overnight using destaining solution (6% acetic acid, 40% methanol in nanopure water). The gel was imaged using a ChemiDoc™ MP Imaging System (Bio-Rad, Hercules, CA, USA).

For SDS-PAGE analysis to evaluate extraction efficiency from different amounts of tissue, samples were diluted with an equal volume of 2X Laemmli Sample Buffer. 1 µL PageRuler™ unstained protein ladder (Thermo Scientific) or 1 µg total protein per sample was loaded into individual wells of a hand-cast 12.5% polyacrylamide SDS-PAGE gel (15 comb well, 10 cm × 10 cm, 1 mm thick). Samples were electrophoretically migrated through the stacking gel at 70 V for 30 min, then through the resolving gel at 150 V for 90 min. The gel was washed briefly with nanopure water, fixed in fixing solution (50% methanol, 7% acetic acid in nanopure water) for 30 min, and then stained with SYPRO™ Ruby gel stain (Thermo Scientific) overnight. After overnight staining, the gel was washed in washing solution (10% methanol, 7% acetic acid in nanopure water) for 30 min. The gel was imaged using a ChemiDoc™ MP Imaging System (Bio-Rad).

**Bottom-Up Data Acquisition without TIMS/PASEF.** For experiments collected with TIMS and PASEF turned off, 200 ng of K562 peptide digest was loaded on a capillary C18 column (25 cm length, 75 µm inner diameter, 1.6 µm particle size, 120 Å pore size; IonOpticks, Fitzroy, VIC, AUS). Peptides were separated at 55 °C using a 120 min gradient at a flow rate of 400 nL/min (mobile phase A (MPA): 0.1% FA; mobile phase B (MPB): 0.1% FA in acetonitrile). A stepwise gradient of 2-17% MPB was applied for 60 min, followed by a step from 17-25% MPB from 60-90 min, 25-37% MPB from 90-100 min, 37-85% MPB from 100-110 min, and finished with a wash at 85% MPB for an additional 10 min for a total runtime of 120 min per analysis, and peptides were eluted from the column into the timsTOF Pro (Bruker Daltonics, Bremen, Germany). Mass spectra for MS and MS/MS scans were recorded between 100-1700 m/z. MS scans were collected at 2 Hz and the top 30 most intense ions were selected for collision-induced dissociation using a 3 *m/z* window and a voltage energy scaled by *m/z* and charge, as previously described.[1]

**Data Analysis for K562 Whole-Cell Digest Comparison.** Raw files produced for the comparison of timsOFF and timsON runs of K562 lysate were processed with FragPipe (version 15.0) and MSFragger (version 3.2).[2] MS/MS spectra were matched to the Uniprot human database UP000005640 (accessed 15 April 2021). Searches were performed without quantification using the default parameters. Total number of unique protein groups and unique peptides among three runs for each condition were tabulated and reported in Supplementary Figure S1.

### Supplementary Figures

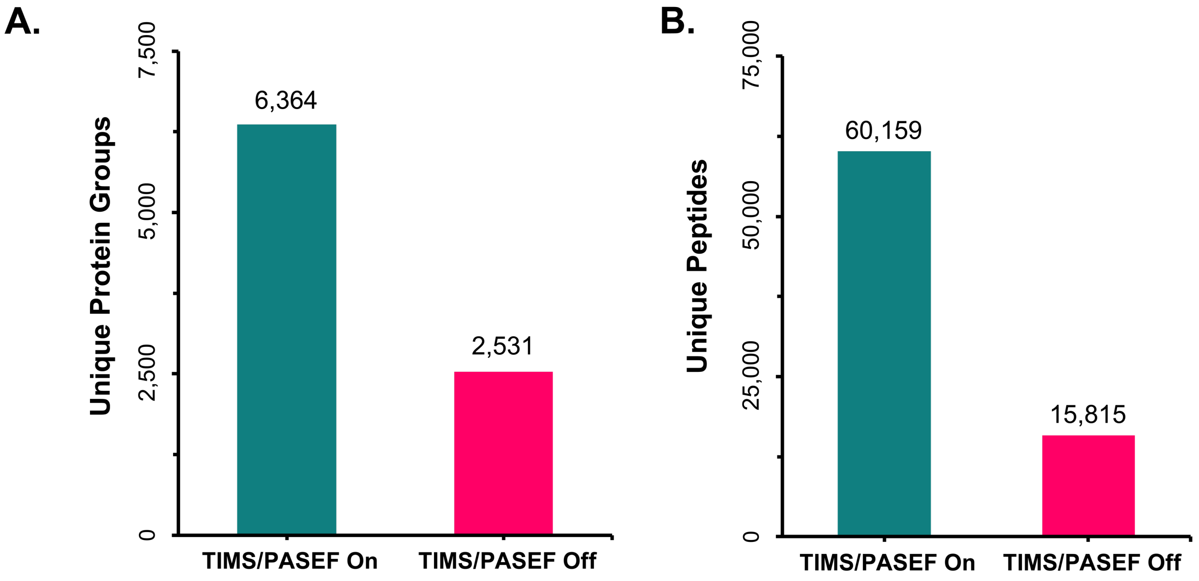


**Supplementary Figure S1**

**timsTOF Pro and PASEF enable deep proteome coverage. (A and B).** Triplicate injections of 200 ng K562 whole cell lysate with TIMS and PASEF either enabled or disabled, showing the summed unique protein groups **(A)** and unique peptides **(B)** identified by MSFragger.[2] Total number of unique identifications from the triplicate injections is displayed above each bar.

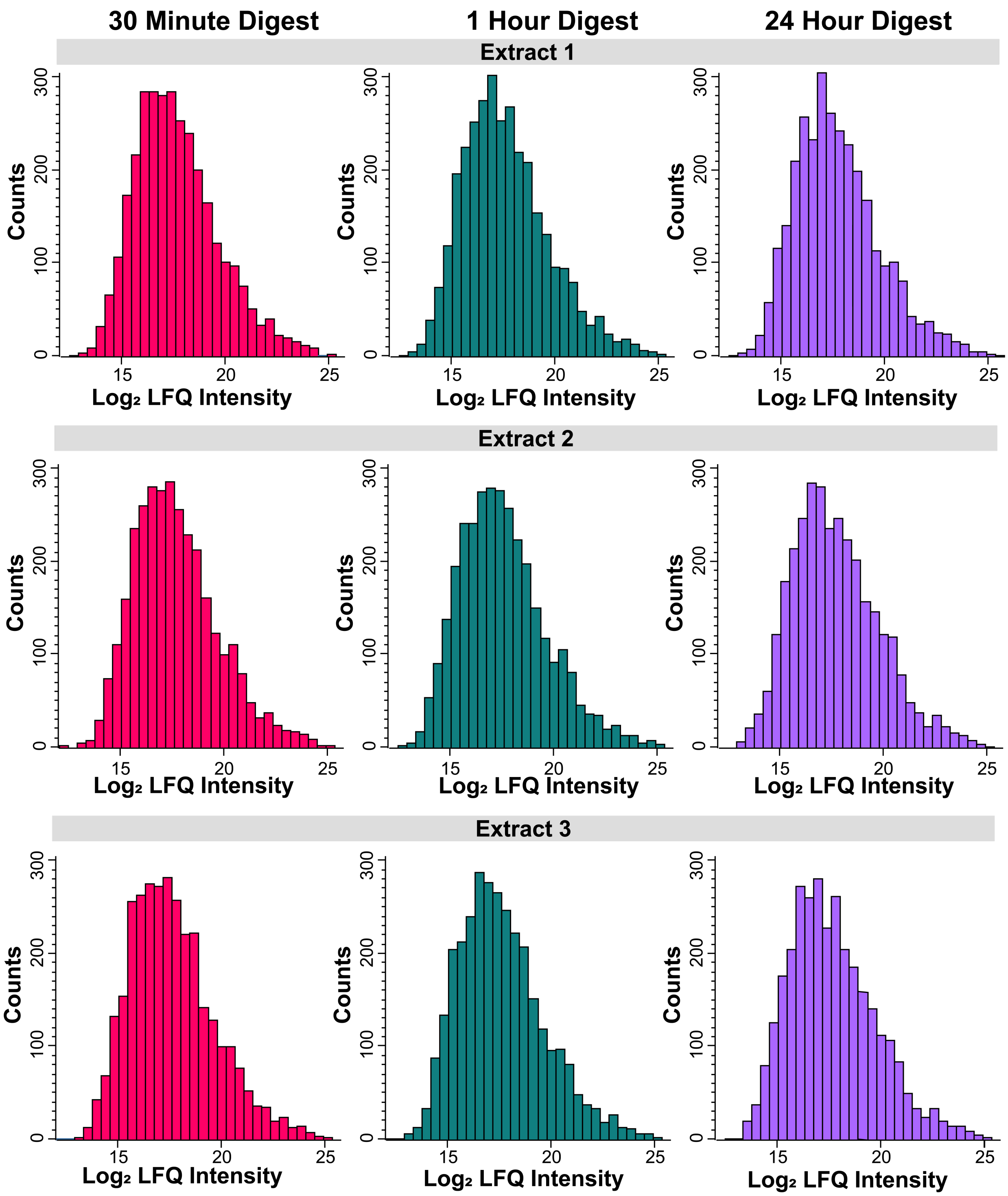


**Supplementary Figure S2.**

**Log$_2$ LFQ protein group intensities for each digestion time are normally distributed.** Histograms of Log$_2$ LFQ protein group intensities displaying a normal unimodal distribution for 30 min, 1 h, and 24 h tryptic digestions in 0.1% Azo for each extraction replicate.

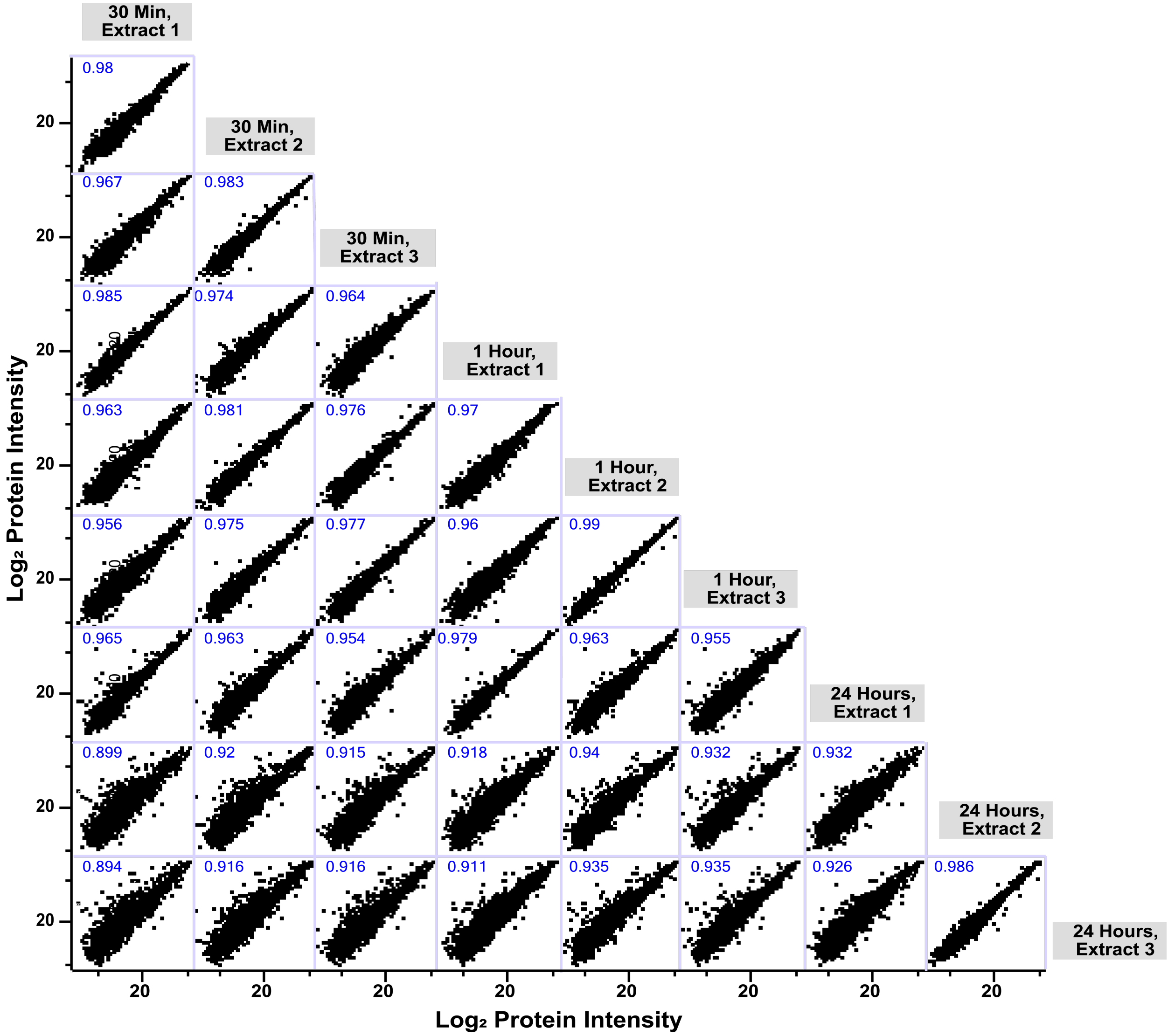


**Supplementary Figure S3.**

**Individual digestion replicates show reproducible $Log_2$ LFQ protein intensities.** Multiscatter array plots of all shared $Log_2$ LFQ protein intensities against one another demonstrating reproducibility within and between digestion groups. Pearson correlation coefficients are displayed in the top left hand portion of each comparison.

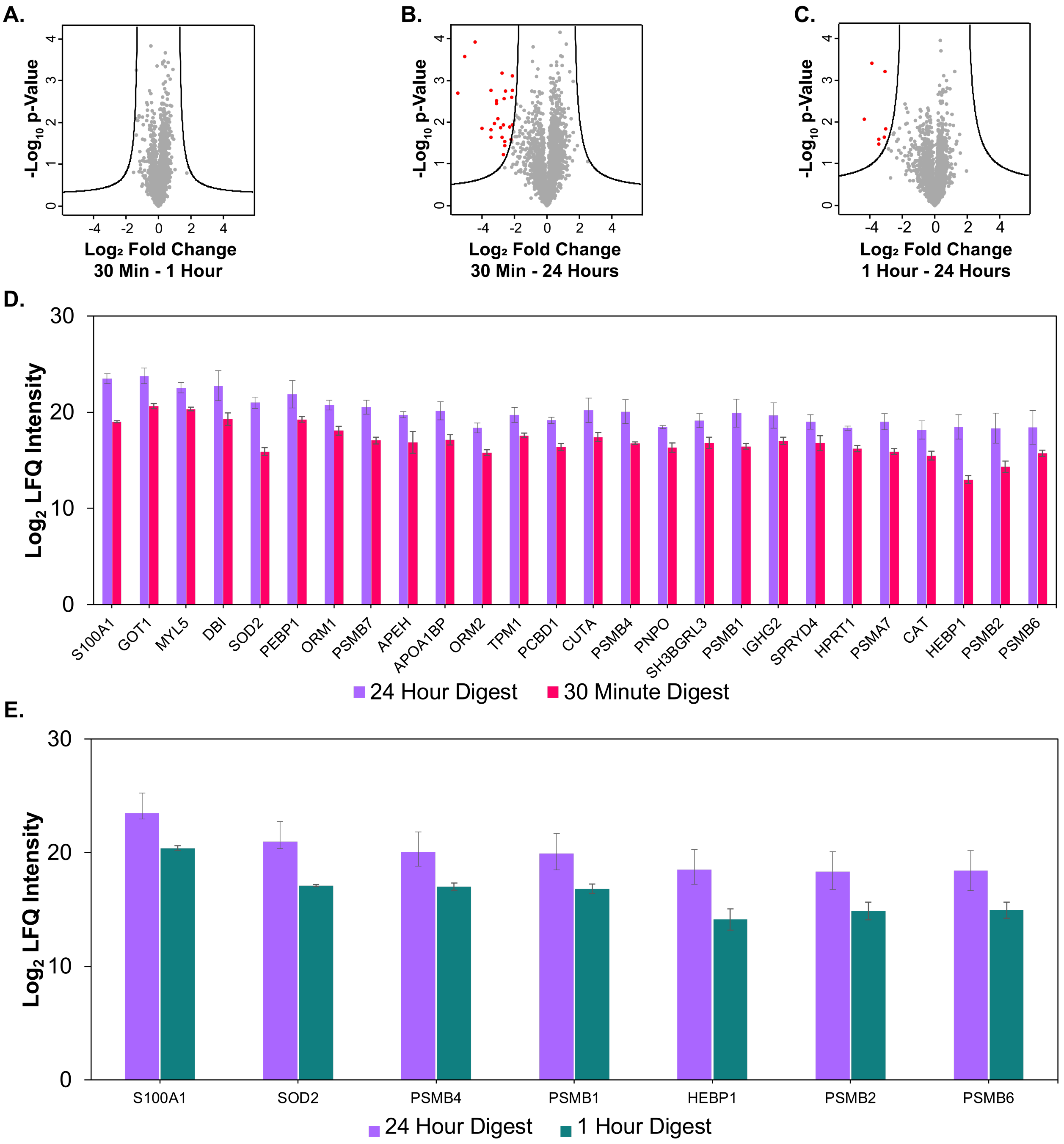


**Supplementary Figure S4.**

**Comparison of protein abundance between different tryptic digestion times. (A-C)**. Proteins with $Log_2$ fold-change≥2 and an adjusted p-value≤0.01 between 30 min and 1 h **(A)**, 30 min and 24 h **(B),** and 1 h and 24 h **(C)** tryptic digestion times. **(D and E).** Bar chart displaying proteins determined to be differentially enriched between 30 min and 24 h **(D)** and between 1 h and 24 h **(E)** tryptic digestions. Bars indicate the average $Log_2$ LFQ intensity between n=3 replicates. Error bars represent SEM.

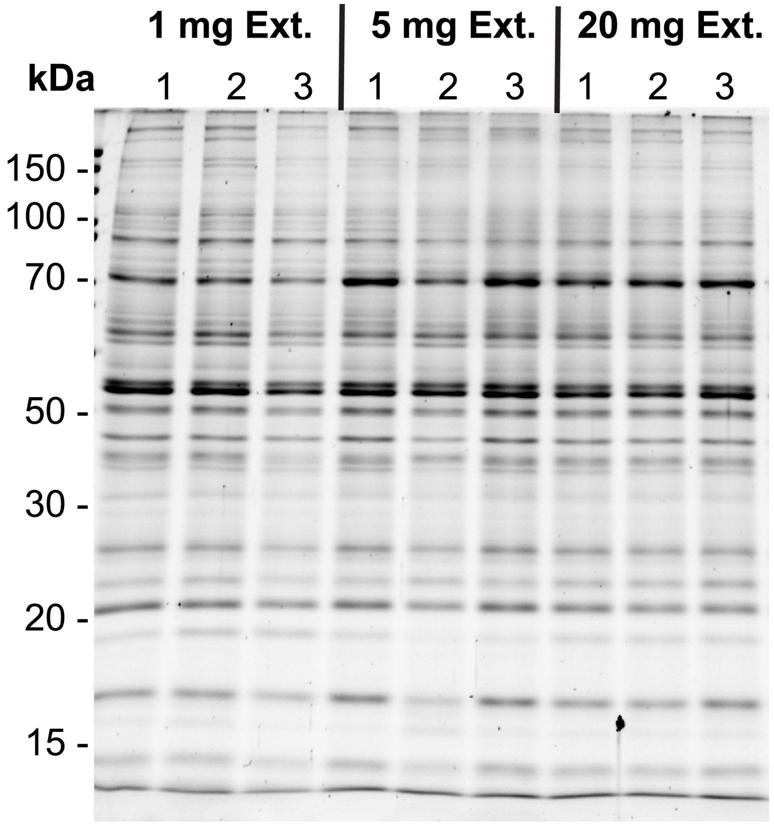


**Supplementary Figure S5.**

**SDS-PAGE demonstrating reproducible proteome extraction from different tissue amounts.** 1 $\mu$g total protein from 3 extraction replicates from 1 mg, 5 mg and 20 mg of tissue separated by 12.5% SDS-PAGE and stained with SYPRO Ruby.

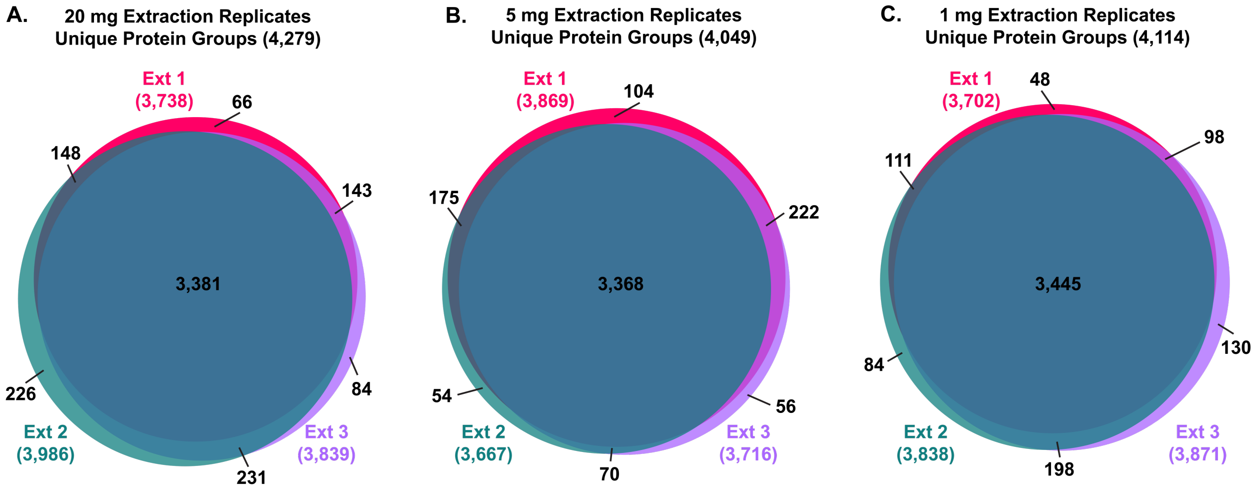


**Supplementary Figure S6.**

**Comparison of protein groups identified between different tissue extraction replicates. (A-C).** Overlap of protein groups identified by MS/MS between extraction replicates from 20 mg **(A)**, 5 mg **(B)**, and 1 mg **(C)** left ventricular (LV) heart tissue extractions after 30 m tryptic digestions.

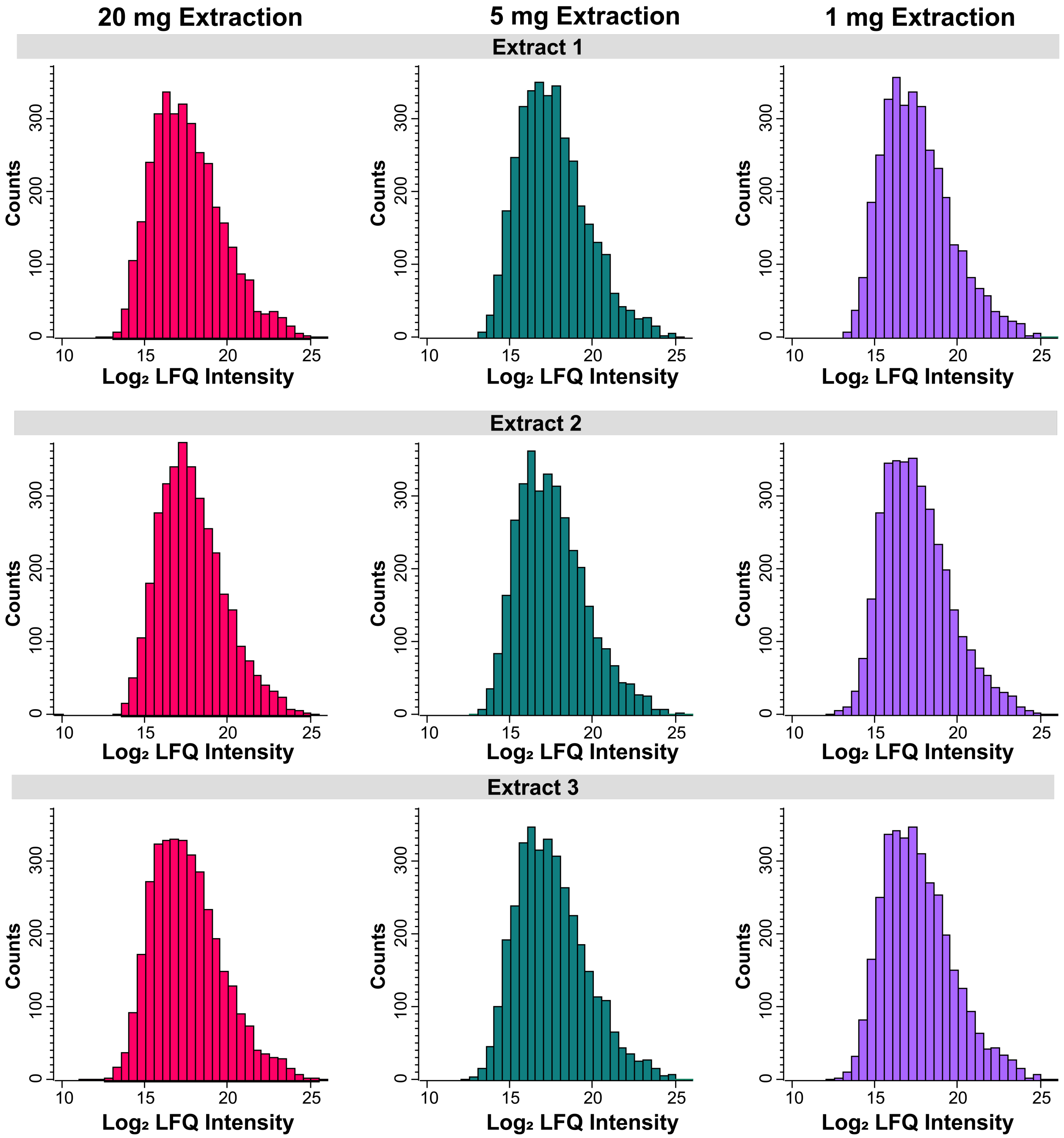


**Supplementary Figure S7.**

**$Log_2$ LFQ protein group intensities for each tissue amount are normally distributed.** Histograms of $Log_2$ transformed LFQ protein group intensities displaying a normal unimodal distribution for 20 mg, 5 mg, and 1 mg tissue extractions (n=3).

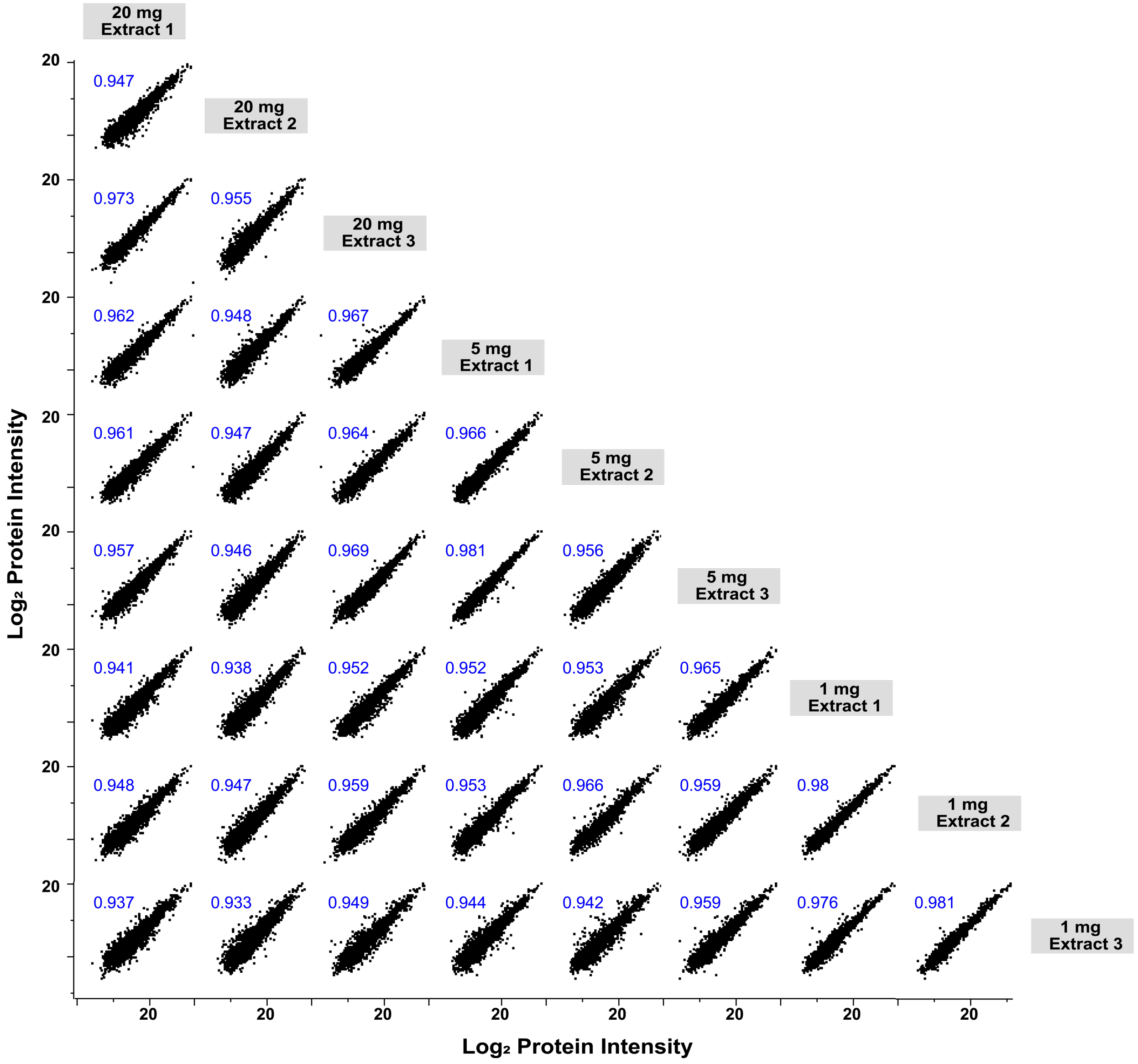


**Supplementary Figure S8.**

**Individual replicates from different tissue amounts show reproducible $Log_2$ LFQ protein intensities.** Multiscatter array plots of all shared $Log_2$ LFQ protein intensities against one another demonstrating reproducibility within and between tissue extraction amounts. Pearson correlation coefficients are displayed in the top left hand portion of each comparison.

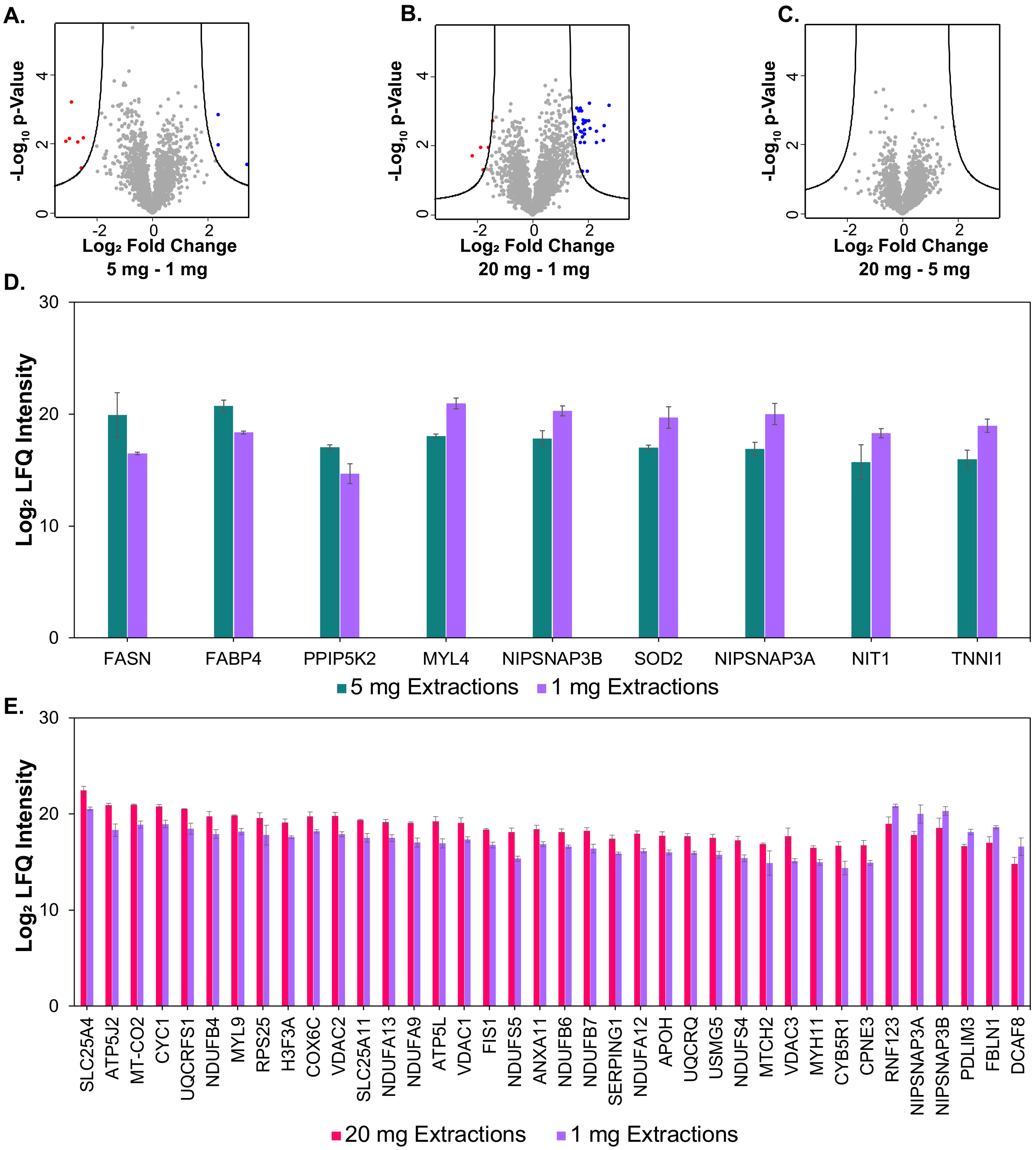


**Supplementary Figure S9.**

**Comparison of proteins extracted between different tissue amounts. (A-C)**. Differentially extracted proteins with $Log_2$ fold-change≥2 and an adjusted p-value≤0.01 between 5 mg and 1 mg **(A)**, 20 mg and 1 mg **(B),** and 20 mg and 5 mg **(C)** tissue extractions. **(D and E).** Bar chart displaying proteins determined to be differentially extracted between 5 mg and 1 mg **(D),** and between 20 mg and 1 mg **(E)** tissue amounts. Bars indicate the average $Log_2$ LFQ intensity between n=3 replicates. Error bars represent SEM.

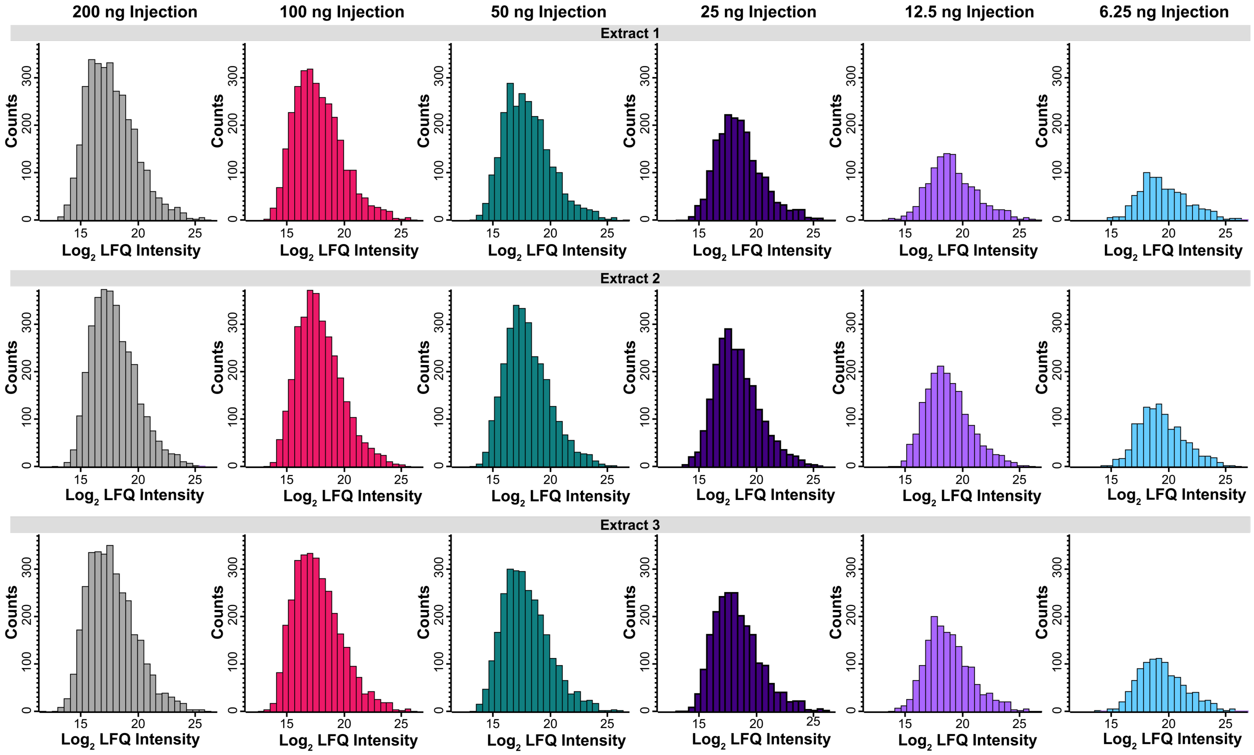


**Supplementary Figure S10.**

**$Log_2$ LFQ protein group intensities are normally distributed at low peptide loading amounts.** Histograms of $Log_2$ transformed LFQ protein group intensities from 200, 100, 50, 25, 12.5 and 6.25 ng injections of total peptide from 20 mg tissue extractions displaying a normal unimodal distribution (n=3).

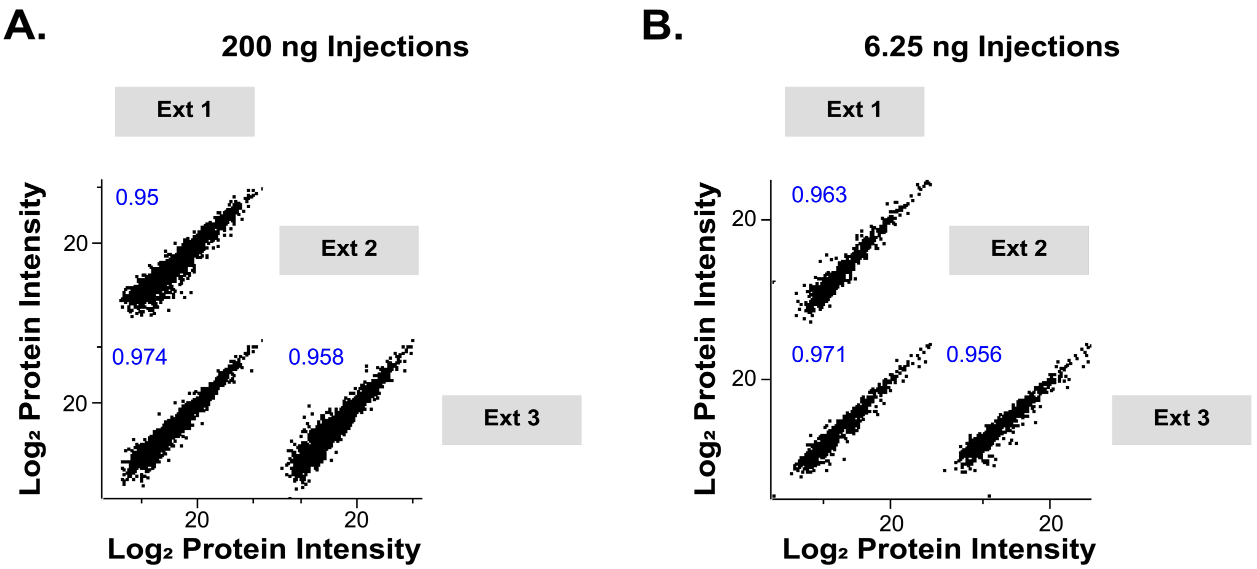


**Supplementary Figure S11.**

**200 ng and 6.25 ng peptide injection replicates show reproducible $Log_2$ LFQ protein intensities.** Multiscatter array plotting all shared $Log_2$ LFQ protein intensities between 200 ng peptide injections **(A)** and 6.25 ng peptide injections **(B)** demonstrating reproducibility for quantitative proteomics even when analyzing low amounts of peptide. Pearson correlation coefficients are displayed in the top left hand portion of each comparison.
